# Proteomic responses of oat (*Avena sativa* L.) to drought stress

**DOI:** 10.1101/2024.09.21.614288

**Authors:** Caijin Chen, Mingfang Bao, Yanxia Zeng, Xuemin Wang, Wenhui Liu

## Abstract

Drought is a major abiotic factor limiting the growth and development of the oat industry, and understanding its drought tolerance mechanisms is vital to oat production. In this study, we measured the phenotypic and physiological indices of drought-resistant (Grain King [G]) and water-sensitive (XiYue [X]) oat varieties and performed comparative proteomic analysis under drought stress and normal water supply (soil water content of 75% ± 5% of field water holding capacity) conditions. The results indicated that plant height, aboveground biomass, and underground biomass of variety X were 7.9%, 9.5%, and 14.6% lower than those under normal water supply, respectively, and the difference in plant height was significant (*p* < 0.05), whereas the decrease in all these indicators of variety G was small. Drought stress significantly increased malondialdehyde (MDA) content, soluble sugar (SS) content, superoxide dismutase (SOD) activity, and peroxidase (POD) activity of variety G by 48.6%, 68.5%, 81.3%, and 101.7%, respectively (*p* < 0.05). Variety X also showed increases up to various extents, but the increases were smaller than those of variety G. Additionally, 151 and 792 differentially expressed proteins (DEPs) identified in varieties G and X, respectively. Weighted Gene Co-expression Network Analysis(WGCNA), Gene Ontology(GO) and Kyoto Encyclopedia of Genes and Genome(KEGG) analyses demonstrated that the DEPs who were highly correlated with POD and SOD activity and SS content in variety G, were majorly involved in energy metabolism, protein translation, RNA processing, amino acid metabolism, and protein folding, whereas those with high correlation with the above three physiological indicators in variety X were primarily involved in RNA processing, protein stabilization, plant photosynthesis, intracellular signal transduction, and protein folding. Overall, the study elucidated the drought resistance mechanisms of different types of oats at the protein level。

## Introduction

With frequent global climate extremes and decreasing availability of irrigation water, drought has become an essential abiotic stressor affecting crop production worldwide [1]. Based on statistics, 43% of arable land worldwide is affected by drought stress [2].serious drought events will continue to increase, especially under climate change scenarios that threaten food production [3]. The damage to crops caused by soil drought decreases crop yield and quality and limits the latitude of crops and the soil environment in which they can be grown [4,5]. In a word, drought is a prominent threat to agriculture worldwide more than ever [6]. However, when crops experience such water deficits for long periods, they evolve complex mechanisms to sense water availability in the environment. They activate stress responses at the morphological, physiological, biochemical, and molecular levels of the plant to reprogram their metabolism and growth to establish new homeostasis [7–9]. Among them, in terms of morphology, most plants respond to drought stress through adaptive changes in root stem and leaf morphology structure, changes in leaf orientation, stomatal closure, deepening of root systems, decrease in branch growth, and leaf senescence and dormancy, etc.[4,8–10]. In terms of physiology and biochemistry, drought stress causes the production and accumulation of organic solutes, including proline, betaine, sugar, and sugar alcohols, etc.To prevent membrane disintegration and enzyme inactivation in plants [11–13]. At the molecular level, drought stress activates multiple drought-related functional genes and induces the synthesis and expression of stress effectors and regulatory proteins, and these transcripts and proteins that mediate drought stress response are primarily involved in transcription, regulation of reactive oxygen species scavengers and antioxidants, and activation of signal transduction pathways, etc.[14–15].Thus, plants enhance their ability to avoid damage (stress avoidance mechanism) or maintain their metabolic functions under drought conditions (stress tolerance mechanism) [16].

Oat (*Avena sativa* L.) is one of the six largest cereal crops worldwide, with more than 10 million hectares under cultivation worldwide, and is mainly used as food, feed, and forage to grow [16,17]. Oat contains highly valuable compounds for industrial applications, such as glucan, oil, and protein [18,19], which lower blood cholesterol levels and minimize the risk of heart disease [20], and in recent years, oat has become one of the most loved crops globally. In China, with the implementation of a series of policies such as “grain to feed” and the rapid development of animal husbandry, the demand for high-quality forage oats has increased significantly, and planting areas have been expanding. However, drought has seriously affected the productivity of oats, and studies have substantiated that in most countries where oats were grown, they were commonly grown in drought-prone or nutrient-poor areas used for the production of major food crops such as wheat [21]. Therefore, it is imperative to conduct research on drought-related genes and protein in oats to analyze the drought resistance mechanism of forage oat at the molecular level to better guide the selection and breeding of drought-resistant varieties of forage oats, thereby steering their production.

Proteomics is the study of the function, recognition, and regulation of a complete set of proteins in a cell, subcellular compartment, and tissue. It is essential to our understanding of the complex biological processes of drought stress at the molecular level. Many proteomic studies have elucidated the effects of drought stress on plant growth and development and the protein expression strategies of various plants in response to drought stress. For example, drought was shown to result in a decrease in the content of rubisco-binding proteins and inhibited photosynthesis in alfalfa leaves [22]. Drought was also shown to upregulate protective and stress-related proteins (mainly chaperone proteins and dehydrins) in two genotypes of maize-sensitive and -tolerant varieties, but changes in antioxidant enzyme activities and differences in the levels of various detoxification proteins correspond one-to-one [23]. In another study, drought stress resulted in a decrease in photosynthesis, carbohydrate levels, protein levels, and defense- and energy metabolism-related protein levels in safflower drought-tolerant variety and an increase upon rehydration, but sensitive variety did not differ under drought and rehydration conditions [24]. Proteins involved in osmoregulation, defense signaling, and programmed cell death in soybean roots that are expressed in response to short-term drought stress are critical in drought adaptation in soybean plants [25]. Additionally, a study reported that under mild drought stress, osmotic stress cells (such as cottonseed sugars and proline) accumulated and improved resistance, while under moderate and severe drought stress, the oxidation of unsaturated fatty acids and accumulation of glucose and galactose increased, but the synthesis of β-coumarin and the terpene precursor 2,3-epoxide was suppressed and the accumulation of triterpenes (glycyrrhetinic acid) was affected [26]. Thus, proteomic studies on plant response to drought are necessary to understand the complex regulatory mechanisms of plant drought resistance. However, the mechanism of protein response to drought stress in forage oats remains unclear. Therefore, in this study, we used tandem mass tag (TMT) chemical marker quantitative proteomics technology to characterize and quantify proteins in the seedling leaves of various drought-resistant forage oat varieties under drought stress and normal water supply treatment. Additionally, we screened differential protein expression profiles using comparative analysis to discover the key proteins that play essential roles in drought stress stimulation and regulation pathways in forage oats as well as to preliminarily resolve their functions and mechanisms. Our findings will serve as a foundation for further research on the mechanisms of oat response to drought stress.

## Materials and methods

### Plant materials

The materials used in this study were the commercial varieties of forage oats, A. *sativa cv*. Grain King (provided by Baili International Grass, Beijing Co., Ltd.) and A. *sativa cv.* XiYue (provided by Ningxia Daxinong Seed Co.). In a previous study, the phenotypic traits and survival rates of 13 oat varieties were determined under drought stress and rehydration conditions, and varieties with different drought tolerance levels were screened, of which the variety G was drought-resistant and the variety X was water-sensitive.

### Stress treatment

Two treatments, namely, drought stress and normal water supply, were set up for each variety, with three replicates planned for each treatment. In the drought stress treatment, normal irrigation was implemented from sowing to 15 days after seedling emergence, maintaining the soil water content at 75% ± 5% of the field water holding capacity. After 15 days of seedling emergence, the water was allowed to deplete naturally, and the morphology indicators was measured after 7 days of stress, while the penultimate three leaves of single plants were selected and snap-frozen in liquid nitrogen and stored in a refrigerator at

−80°C for the measurement of physiological indexes and proteomics analysis. In the normal water supply treatment (the control), the soil water content was maintained at 75% ± 5% of the field water holding capacity until sampling. The sampling time, sampling technique, and measurement index were the same as those of the drought stress treatment. Water control was performed by the weighing method, and watering was done once a day to ensure that the water supply met the requirements of the experiment.

### Plant growth conditions

Each pot was tested, and soil water content, field water holding capacity, and relative soil water content were calculated. Based on the data obtained, the pot weight was calculated for each treatment when the set relative water content was achieved. Subsequently, impurities from the seeds of the two varieties were eliminated, and 240 seeds (1 pot for each replicate planting, 80 plants per pot) that were full, uniform, and free of defects and diseases were each selected and washed thrice with distilled water, disinfected with 5% Na_2_ClO_3_, and washed five times with distilled water, dried, and sown in pots (height, 24.5 cm; outer diameter at the top, 35 cm; inner diameter, 30 cm; inner diameter at the bottom, 20 cm). The pots were finally moved to controlled growth room. The potting soil was obtained from the mixture of soil at 0-30 cm of the cultivation layer and substrate at the experimental site of Qinghai University experimental field, and the basic physical and chemical qualities of the soil are presented in Table 1. The oat seeding were cultured under the following growth conditions:20/25℃(night/day), 50-60% relative humidity, and a photoperiod of 8h/16h(night/day) by 500 μmol m^-2^.s^-1^ light intensity. The potted plants moved into the controlled growth room were manually interplanted when they reached three leaves keeping 40 healthy plants in each pot eventually.

**Table 1.** Basic physical and chemical properties of soils.

| Organic matter (g/kg) | Total nitrogen (mg/kg) | Total phosphorus (%) | Total potassium (%) | Hydrolytic nitrogen (mg/kg) | Fast acting potassium (mg/kg) | Effective phosphorus (mg/kg) | Water-soluble salt Total amount (g/kg) | pH |
| --- | --- | --- | --- | --- | --- | --- | --- | --- |
| 89.33 | 3.42 × 10<sup>3</sup> | 0.17 | 3.00 | 241.08 | 1147.00 | 273.74 | 4.78 | 7.75 |

### Morphological and physiological index determination methods

The morphological indices were determined by referring to the method described by Wenqin Ji [27]. In each treatment, 15 plants were selected for the determination of plant height, and single plants at the end of height determination were sampled, following which their root system was rinsed using distilled water, the aboveground part and the root system were separated and dried, and all of them were placed in an oven at 105℃ ± 2℃ to be killed for 30 min. The samples were then dried at 80℃ ± 2℃ to a constant weight, and the aboveground part and root system of each plant were weighed separately to determine the aboveground biomass and belowground biomass, respectively.

For the measurement of physiological indicators, we determined MDA content by referring to the colorimetric technique of using thiobarbituric acid as described by Na Zhang [28]. Additionally, we determined the content of SS by referring to the anthrone method described by Jun Wang [29]. Furthermore, we determined the total SOD and POD activity using the SOD (A001-3-2) and POD (A084- 3-1) test kits (Nanjing Jiancheng Institute of Biological Engineering, Nanjing, China) using the Water-Soluble Tetrazolium-1 (WST-1) and colorimetric methods, respectively.

### Proteomics analysis Sample preparation

The leaves of samples were first ground to a powder with a small amount of liquid nitrogen, added to 200 µL of lysis buffer (150 mM Tris–HCl, 4% sodium dodecyl sulfate, 100 mM dithiothreitol (DTT), pH 7.8), sonicated, boiled for 5 min, and precipitated with trichloroacetic acid–acetone solution. After centrifugation at 16,000 rpm (16,000 ×g, 1 min, 4°C), the tubes were washed twice with cold acetone and air dried. Each sample tube was added with 150 µL of lysis buffer, sonicated, and centrifuged at 16,000 rpm for 15 min to remove undissolved cell debris. The supernatant was collected for quantification using the bicinchoninic acid protein kit (Bio-Rad, USA).

### Protein digestion

Protein digestion was performed according to the method described by Wisniewski et al. [30]. Briefly, DTT, detergents, and other low-molecular-weight fractions were removed by centrifugal repeat ultrafiltration (Microcon units, 30 kD) using 200 μL of UA buffer (150 mM Tris–HCl, 8 M urea, pH 8.0). Next, 100 μL of 0.05 M iodoacetamide was added to the UA buffer to block the reduced cysteine residues and incubated in the dark for 20 min. The filters were washed thrice with 100 μL of UA buffer and twice with 100 μL of 25 mM NH_4_HCO_3_. Finally, the protein suspension was digested using 4 μg trypsin in 40 μL of 25 mM NH_4_HCO_3_ overnight at 37°C, and the peptides were obtained as a filtrate. The peptide concentration was determined using the nanodrop OD280 method.

### TMT labeling of peptides and fractionation

Peptides were labeled with TMT reagent according to the manufacturer’s (Thermo Fisher Scientific) instructions. To quantify 12 samples, each sample (100 μg peptide equivalent) was reacted with one tube of TMT reagent. After dissolving the sample in 100 μL of 0.05 M Triethylammonium bicarbonate (TEAB) solution at pH 8.5, the TMT reagent was dissolved in 41 μL of anhydrous acetonitrile. The reaction mixture was incubated for 1 h at room temperature, then 8 μL of 5% hydroxylamine to the sample and incubate for 15 minutes to quench the reaction. The 12 Multiplex-labeled samples were pooled together and lyophilized. The TMT-labeled peptide mixture was separated on a Waters XBridge BEH130 column (C18, 2.1 × 150 mm, 3.5 μm) on a Agilent 1290 High Performance Liquid Chromatography (HPLC) operating at 0.3 mL/min. Buffer A consisted of 10 mM ammonium formate and buffer B consisted of 10 mM ammonium formate plus with 90% acetonitrile. Both buffers were adjusted to pH 10 with ammonium hydroxide. A total of 30 fractions of each peptide mixture were obtained and then concatenated to 15 fractions. The fractions were dried and subjected to Liquid Chromatograph-Mass Spectrometer (LC-MS) analysis.

### LC-MS analysis

An appropriate amount of peptide from each sample was separated via chromatography using a nanoliter flow rate Easy nLC1200 chromatography system (Thermo Scientific). The buffers used in the analysis were solution A (0.1% formic acid aqueous solution) and solution B (0.1% formic acid, acetonitrile, and water mixture (where acetonitrile was 95%)). The column was first equilibrated using 100% A liquid. The sample was loaded onto a Trap Column (100 µm × 20 mm, C18, 5 µm, Dr. Maisch GmbH) and passed through the analytical column (75 µm × 150 mm, C18, 3 µm, Dr. Maisch GmbH) for gradient separation at a flow rate of 300 nL/min. The liquid phase separation gradient settings were as follows: 0–2 min, B liquid linear gradient from 2%–8%; 2–71 min, B liquid linear gradient from 8%–28%; 71– 79 min, B liquid linear gradient from 28%–40%; 79–81 min, B liquid linear gradient from 40%–100 %; 81–90 min, B liquid concentration at 100%. Peptides were analyzed via data-dependent acquisition–mass spectrometry using a Q-Exactive HF-X mass spectrometer (Thermo Scientific) when they had separated after. The analysis time was 90 min, detection mode was positive- ion, parent ion scan range was 400– 1800 m/z, and primary mass spectrometry resolution settings were as follows: 60,000 @m/z 200; (Automatic Gain Contro) AGC target, 3e6; primary Maximumit, 50 ms. secondary mass spectrometry of Peptide segments was collected performed as follows: 20 secondary mass spectra (MS2 scan) of the highest intensity parent ions were triggered after each full scan with the following secondary mass resolution, 45,000 @ m/z 200; AGC target, 1e5; secondary Maximumit, 50 ms; MS2 Activation Type, High-energy Collisional Dissociation (HCD); Isolation window, 1.2-m/z; Normalized collision energy, 32.

### Database query and analysis

The obtained LC–MS/MS raw files were imported into the search engine in Proteome Discower software (version 2.4, Thermo Scientific) for database search. Because there is no oat-specific database, we used the Uniprot-Pooideae database [147368]_924681_20211014.fasta (download date: October 14, 2023) available at the https://www.uniprot.org/taxonomy/147368 Protein Data Bank with 924681 protein sequence. An initial search was set in a precursor to the 6 ppm mass window. The study followed the rules of enzymatic cleavage of trypsin/P, allowing a fragment ion mass tolerance value of 20 ppm and two maximum missing cleavage sites with the following modification sets: fixed modifications, carbamoylmethyl (C), TMT16plex (K), TMT16plex (N term); variable modifications, oxidation (M), acetyl (Protein N term). Peptides require at least six amino acids and one unique peptide per protein. For peptide and protein identification, the false discovery rate (FDR) was set to 1%. TMT reporter ion intensity were used for quantification. Relative quantification of sample proteins was performed using the MaxQuant algorithm (http://www.maxquant.org) [31].

### Bioinformatics analysis

Bioinformatics data were analyzed using the Perseus software, Microsoft Excel, and R statistical software. Differentially expressed proteins (DEPs) were screened with the cutoff of a ratio fold-change of >1.20 or <0.83 and *P*-value of <0.05. Additionally, proteins with significant variance features (FDR) *q* < 0.01) were identified using hierarchical clustering (Euclidean distance) and analysis of variance (ANOVA). For the enrichment analysis of specified protein functions, annotation information was extracted from the UniProtKB/Swiss-Prot, KEGG, and GO databases. GO and KEGG enrichment analyses were carried out with Fisher’s exact test, and FDR correction for multiple testing was also performed. GO terms were grouped into three categories: biological process (BP), molecular function (MF), andcellular component (CC). Correlations between sample features and protein expression changes were analyzed using WGCNA, and expression data were clustered hierarchically based on protein level.

## Results

### Effects of drought stress on the growth of oats

Drought harms the plants, but there are some differences between drought-resistant and water-sensitive varieties. After drought stress, the GD and XD showed different degrees of reduction in plant height, but those of between GD and GW was not significant, whereas the plant height of XD was significantly reduced (by 7.9%) compared with that of XW (*p* < 0.05). Additionally, the aboveground biomass of GD was 9.5% lower than that of GW, while the aboveground biomass of XD was 14.6% lower than that of XW, with no significant differences between treatments within varieties (*p* > 0.05). Furthermore, the underground biomass of GD was 8.0% lower than that of GW, while the belowground biomass of XD was 6.9% lower than that of XW, with no significant differences between treatments within varieties (*p* 0.05) (Figure 1a–c). Taken together, variety G exhibited greater drought resistance than variety X. The drought stress and normal water supply treatments of variety G were indicated as GD and GW, while those of variety X were indicated as XD and XW, respectively.

**Figure 1.**
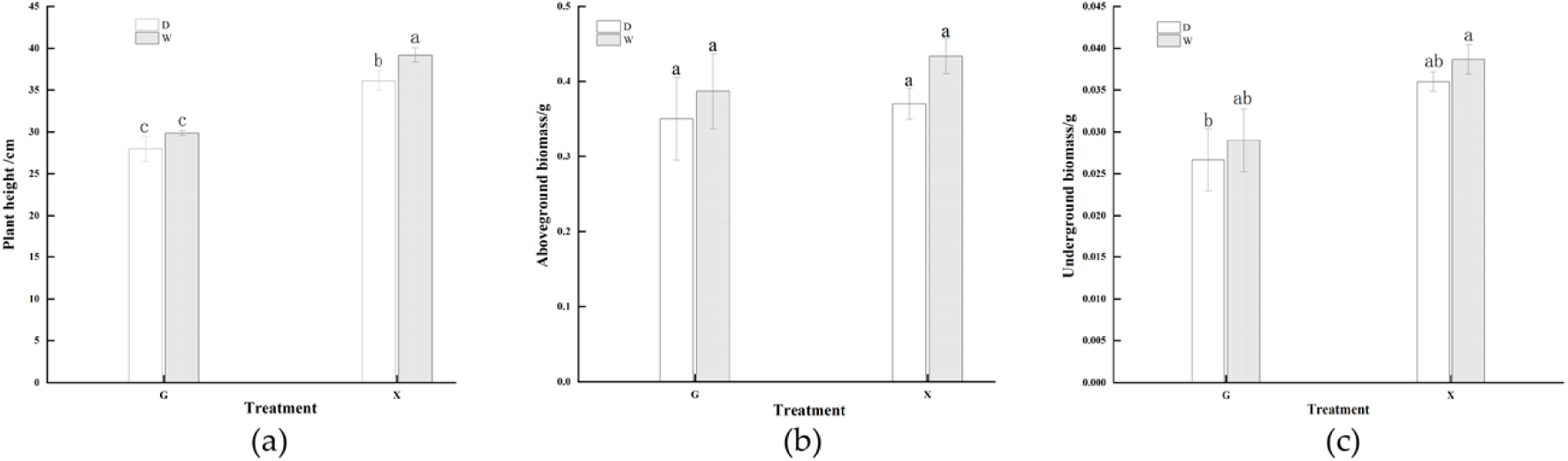
Growth of oat under 7 days of drought stress. (a) Plant height of the two varieties under the two treatments. (b) Aboveground biomass of the two varieties under the two treatments. (c) Belowground biomass of the two varieties under the two treatments. Lowercase letters indicate differences that are significant at p < 0.05.

### Effects of drought stress on the physiological indicators of oats

To further clarify the drought resistance of the two different varieties, many physiological indicators of the plants were measured after drought stress. After drought stress, the MDA content of GD was significantly higher (by 48.6%) than that of GW, while the MDA content of XD was significantly higher (by 28.9%) than that of XW (*p* < 0.05). The SS content of GD was significantly higher (by 68.5%) than that of GW, while the SS content of XD was significantly higher (by 37.2%) than that of XW (*p* < 0.05).

The SOD activity of GD was significantly higher (by 81.3%) than that of GW, while the SOD activity of XD was significantly higher (by 14.6%) than that of XW (*p* < 0.05). The POD activity of GD was significantly higher (by 101.7%) than that of GW, while the POD activity of XD was significantly higher (by 82.5%) than that of XW (*p* < 0.05) (Figure 2). The above analysis suggest that drought stress affects the physiological indicators of oats, and osmoregulatory substances and antioxidant enzyme activities in oats were significantly increased under drought stress. However, there were significant differences between varieties under different treatments, and compared to normal water supply treatment, variety G showed higher change margin in physiological parameters than variety X under drought stress conditions, and stronger drought tolerance, while variety X showed weaker drought tolerance.

**Figure 2.**
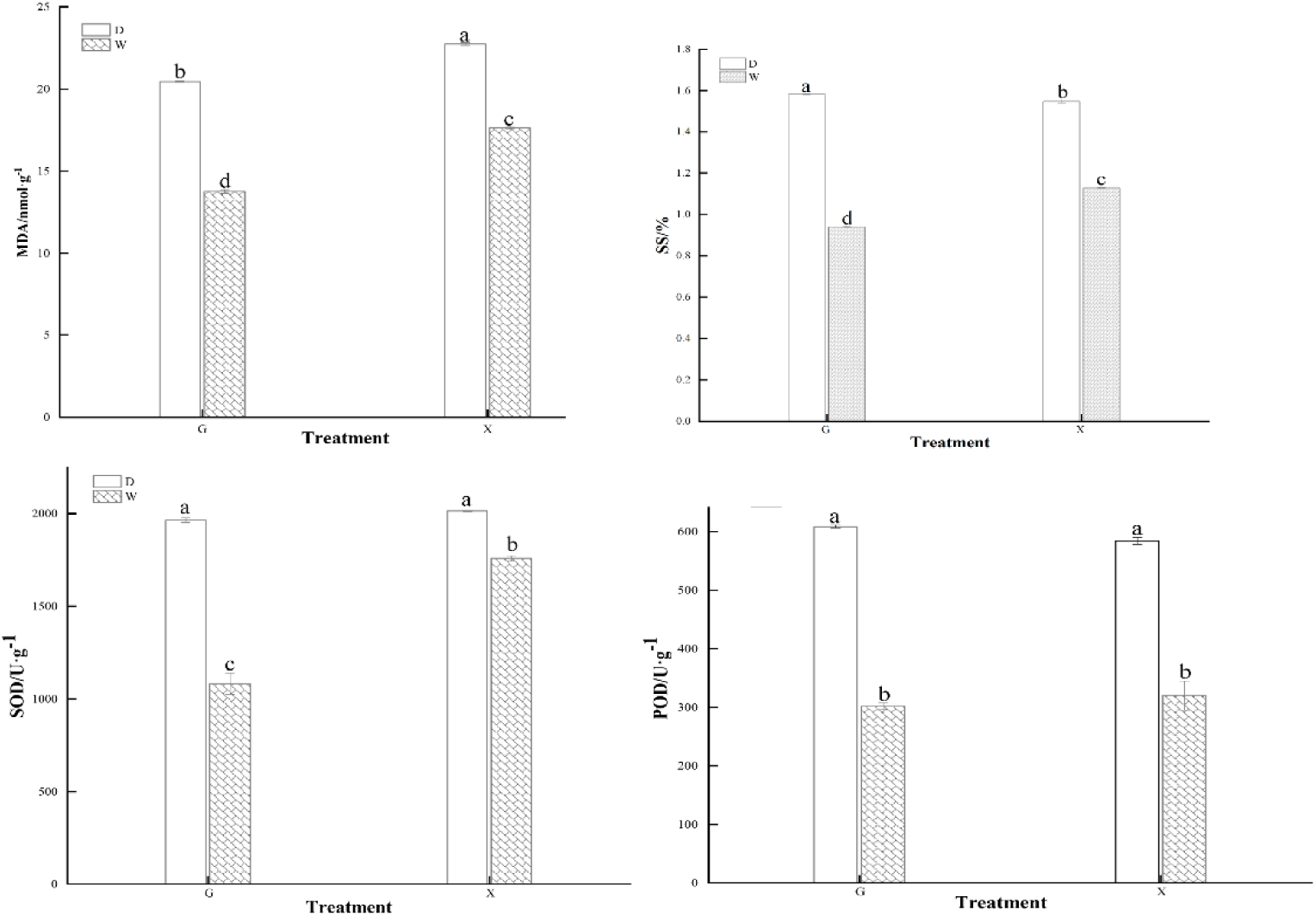
MDA content, SS content, SOD activity, and POD activity of varieties G and X under drought stress and normal water supply treatments.

### Proteomics information identification

The seedling samples of the two oat varieties under different drought stress treatments were identified and the peptide spectral matching (PSM), unique peptide, protein groups, and quantified protein were obtained to be 17690, 4946, 1821, and 1817, respectively. Reproducibility tests of the relative quantification of proteins of the two oat varieties under different treatments showed that the samples were well reproducible, separated between sample groups and close together within groups, and statistically consistent across samples (Figure 3a–c).

**Figure 3.**
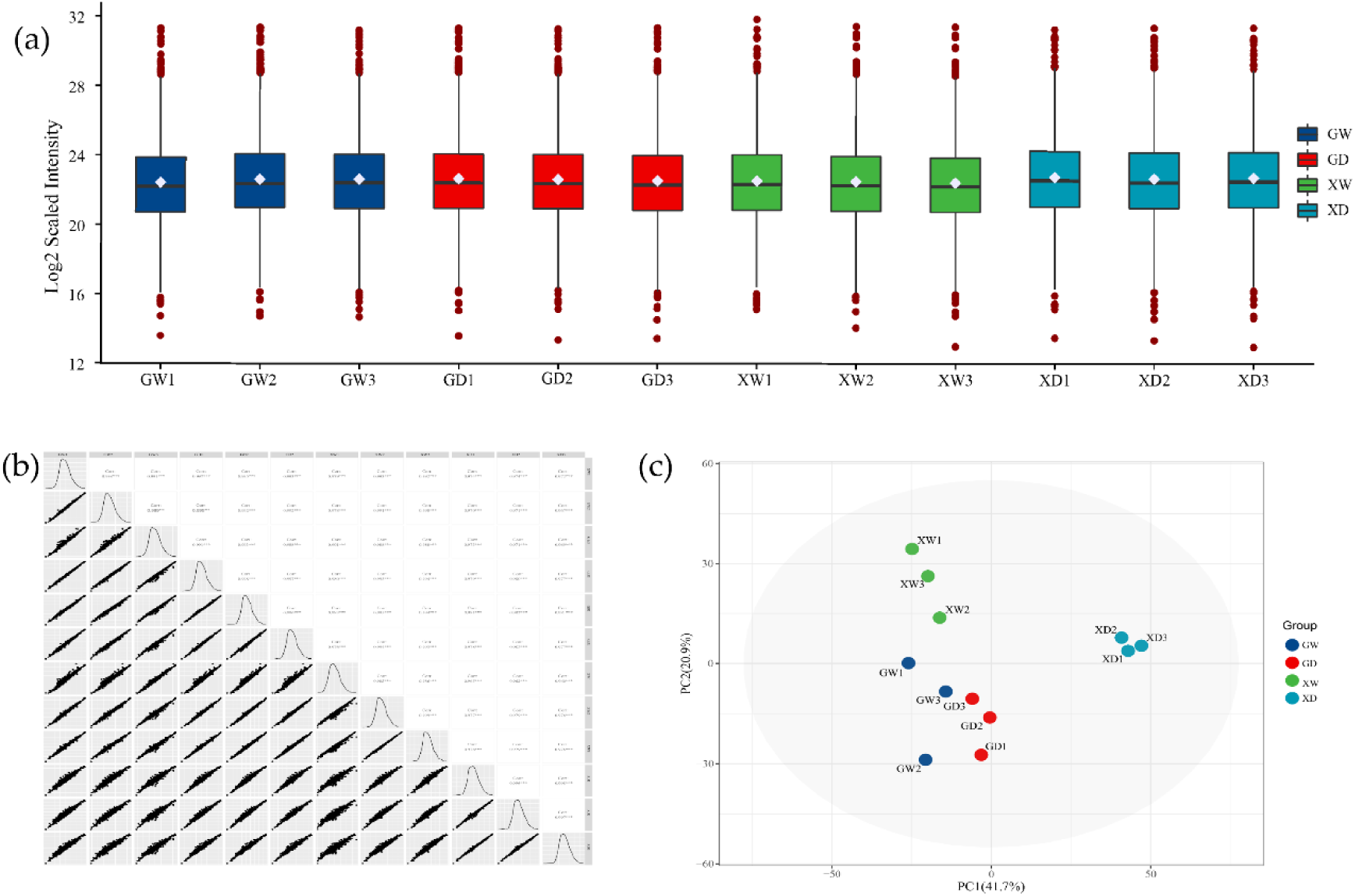
Normalized sample plots for each treatment for data quality control. (a) Normalized density line box plot. (b) Normalized density correlation plot. (c) Normalized principal component plot.

### Analysis of differentially expressed proteins (DEPs)

A total of 846 DEPs (differential expression criteria: p < 0.05, the cutoff of a ratio fold-change of> 1.2 or < 1/1.2) were identified using quantitative analysis of proteins from different treatments of the two drought-resistant varieties. Of these 846 DEPs, 724 were upregulated and 122 were downregulated.

Compared with GW, GD had 140 upregulated DEPs and 11 downregulated DEPs, and compared with XW, XD had 666 upregulated DEPs and 126 downregulated DEPs. These results indicate that there were remarkable differences in protein expression between the two oat varieties under drought stress conditions. The DEPs in the GD vs. GW and XD vs. XW groups are shown in Figure 4a–b.

**Figure 4.**
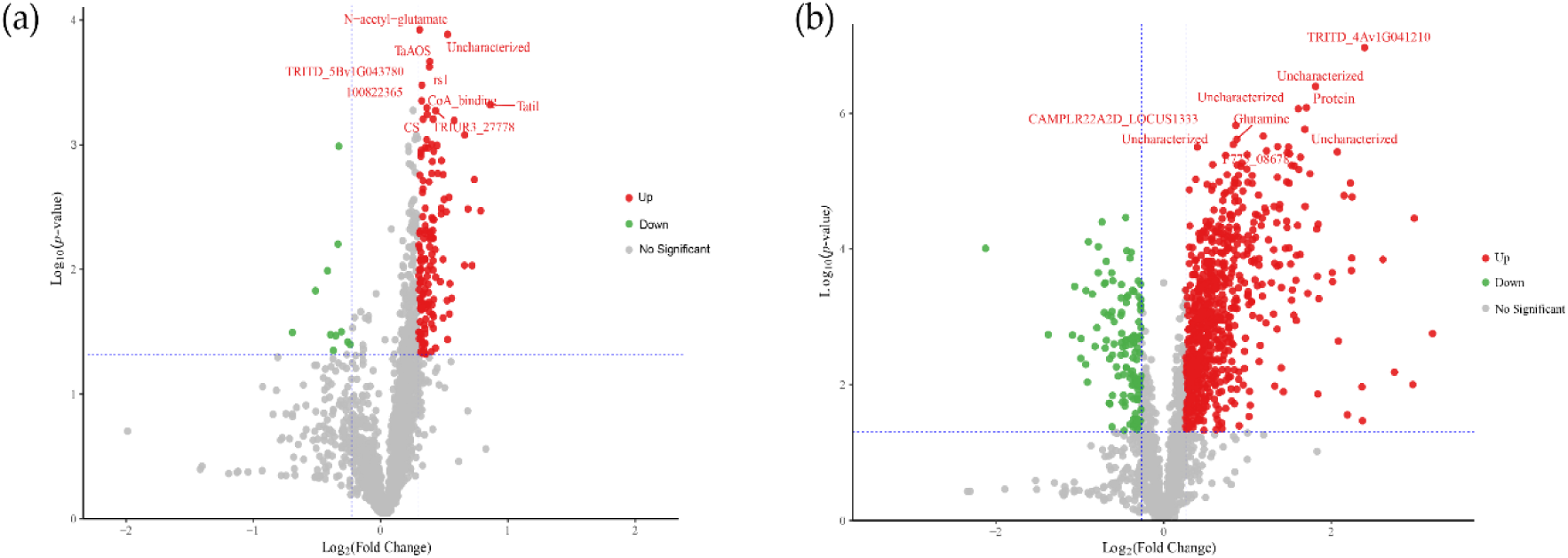
Volcano map of differentially expressed proteins (DEPs). (a) DEPs of the GD vs. GW group. (b) DEPs of the XD vs. XW group. Red indicates significantly upregulated proteins while green indicates significantly downregulated proteins.

### Identification of WGCNA modules related to drought resistance of oat varieties

Nine WGCNA modules were identified using WGCNA of sample characteristics and protein expression changes for the 1817 proteins quantified (Figure 5a–b). The results of the module–trait relationship indicated that 55 proteins in the “MEpink” module were highly correlated with POD (*r* = 0.82, *p* = 0.001) and SOD (*r* = 0.79, *p* = 0.002) activities and SS content (*r* = 0.79, *p* = 0.002), whereas they were moderately correlated with MDA content (*r* = 0.65, *p* = 0.02) and not significantly correlated with plant height (*r* = −0.37, *p* = 0.2), aboveground biomass (*r* = −0.3 *p* = 0.3), and belowground biomass (*r* = −0.2, *p* = 0.5). This may be attributable to a response threshold for drought stress in phenotypic, physiological, and molecular aspects of plant tissues and organs. Within a certain threshold, drought stress only activates the stress response in the physiological and molecular aspects of oats, and the effects on morphological and growth traits are not particularly evident.

**Figure 5.**
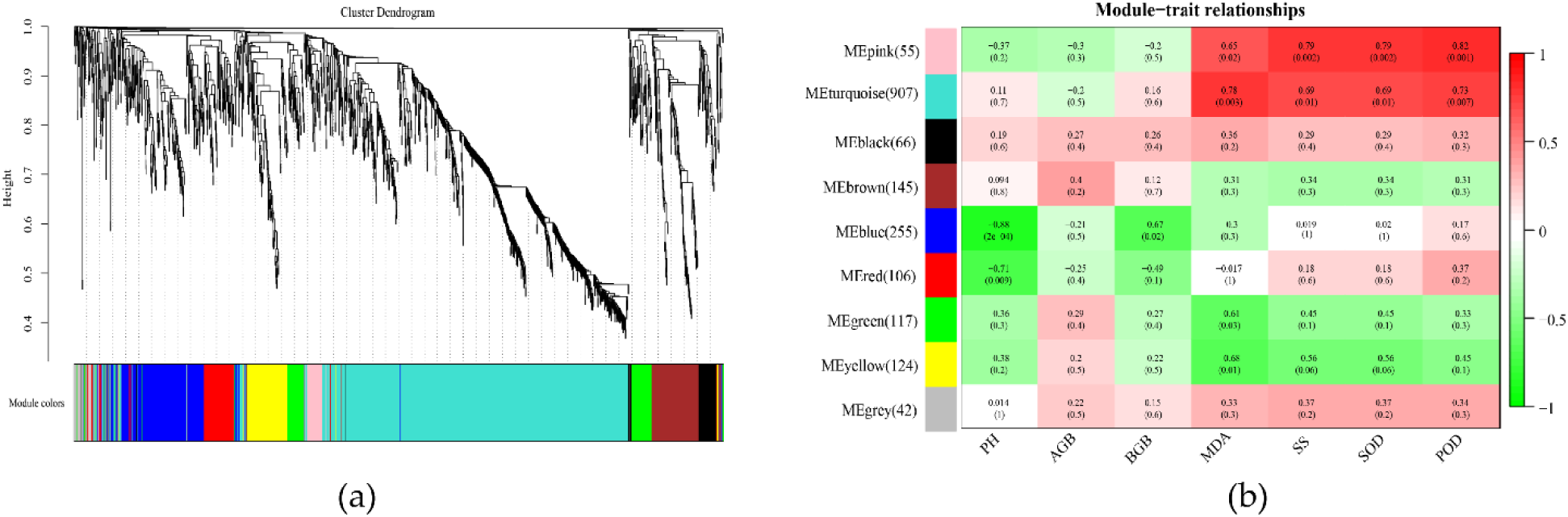
Weighted gene co-expression network analysis (WGCNA) of sample characteristics and proteins of the two varieties with different drought tolerance under drought stress treatment. (a) Stratified clustering of the nine WGCNA modules. (b) Correlation analysis of sample characteris-tics and proteins with changes in expression. Significant differences are indicated in parentheses, correlation coefficients are indicated outside parentheses, PH represents plant height, AGB repre-sents aboveground biomass, and BGB represents belowground biomass.

In the MEpink module, the drought-resistant variety GD had 17 DEPs compared with those of GW, the water-sensitive variety XD had 5 DEPs compared with those of XW, and the two varieties had a total of 2 DEPs, all of which were upregulated. These findings indicate that the 15 DEPs (excluding the two shared ones) in the MEpink module of drought-tolerant variety may play an important role in the drought tolerance of oats, and their synergistic expression can improve drought tolerance.

### GO enrichment analysis of DEPs

GO enrichment analysis of DEPs in the leaves of the two different drought-resistant oat varieties according to GO level two indicated that all DEPs were enriched in biological process (BP), cellular component (CC), and molecular function (MF), and specific proteins with different abundances were classified as shown in Figure 6a–b. In variety G, BP mainly showed enrichment of the following GO terms: cellular process (63, 44.1%), metabolic process (59, 41.3%), response to stimulus (8, 5.6%), regulation of biological processes (3, 2.1%), biological regulation (3, 2.1%), and localization (2, 1.4%), among 11 other aspects. Additionally, CC showed enrichment of only two GO terms: anatomical entity (40, 66.7%) and protein-containing complex (20, 33.3%). Furthermore, MF mainly included the terms catalytic activity (68, 43.6%), binding (68, 43.6%), structural molecule activity (11, 7.1%), and antioxidant activity (4, 2.6%; Figure 6a). In variety X, BP mainly showed enrichment of cellular process (348, 42.7%), metabolic process (323, 39.6%), response to stimulus (45, 5.5%), localization (32, 3.9%), biological regulation (25, 3.1%), and regulation of biological processes (17, 2.1%). Additionally, in CC, only two terms, anatomical entity (156, 63.9%) and protein-containing complex (88, 36.1%), were enriched here as well. Furthermore, the DEPs in MF primarily had 10 functions, including catalytic activity (398, 44.7%), binding (3704, 1.5%), structural molecule activity (37, 4.2%), antioxidant activity (25, 2.8%), translation regulator activity (24, 2.7%), and transporter activity (23, 2.6%). However, the 20 DEPs of the MEpink module were not significantly enriched in the GO database. The above results indicated that DEPs of enrichment levels in terms of CC, MF, and BP were markedly different between the two drought-resistant oat varieties under drought stress conditions, showing that there were large differences in the protein levels of the two varieties in response to drought stress.

**Figure 6.**
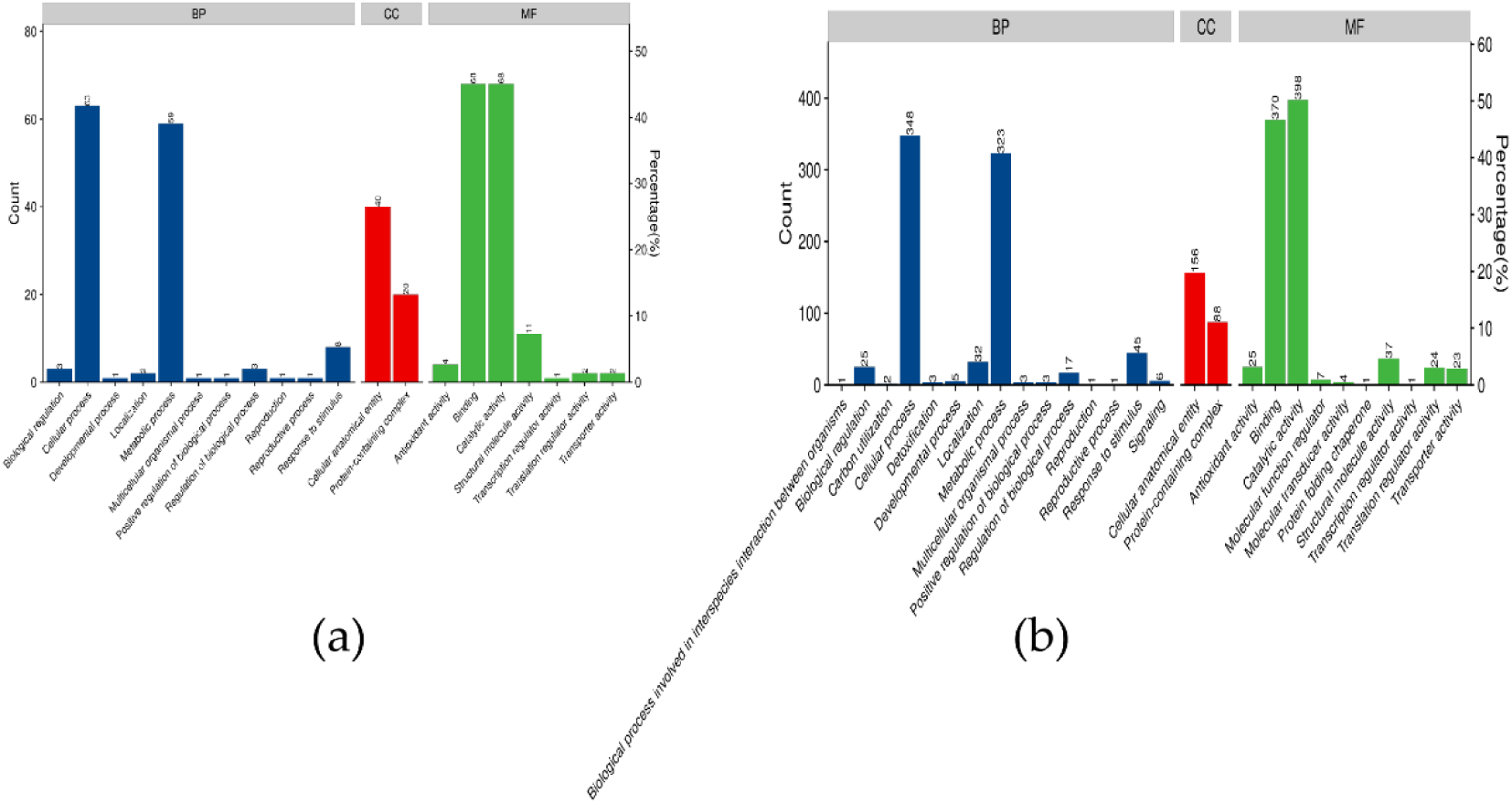
Gene ontology (GO) enrichment of DEPs including three branches: BP, CC, and MF. (a) GO enrichment of DEPs in the GD vs. GW group. (b) GO enrichment of DEPs in the XD vs. XW group.

### KEGG pathway enrichment analysis of DEPs

KEGG pathway enrichment analysis showed that DEPs were enriched in 46 and 97 KEGG metabolic pathways in the G and X varieties, respectively, under drought stress. In variety G, DEPs were significantly enriched in 27 KEGG pathways under drought stress (*p* < 0.05), primarily in metabolic pathways, photosynthesis, nitrogen metabolism, porphyrin and chlorophyll metabolism, oxidative phosphorylation, and protein processing in the endoplasmic reticulum (Figure 7a). In variety X, DEPs were significantly enriched in 51 KEGG pathways under drought stress (*p* < 0.05), primarily in metabolic pathways, carbon metabolism, photosynthesis, biosynthesis of secondary metabolites, glyoxylate and dicarboxylate metabolism, and carbon fixation by photosynthetic organisms (Figure 7b).

**Figure 7.**
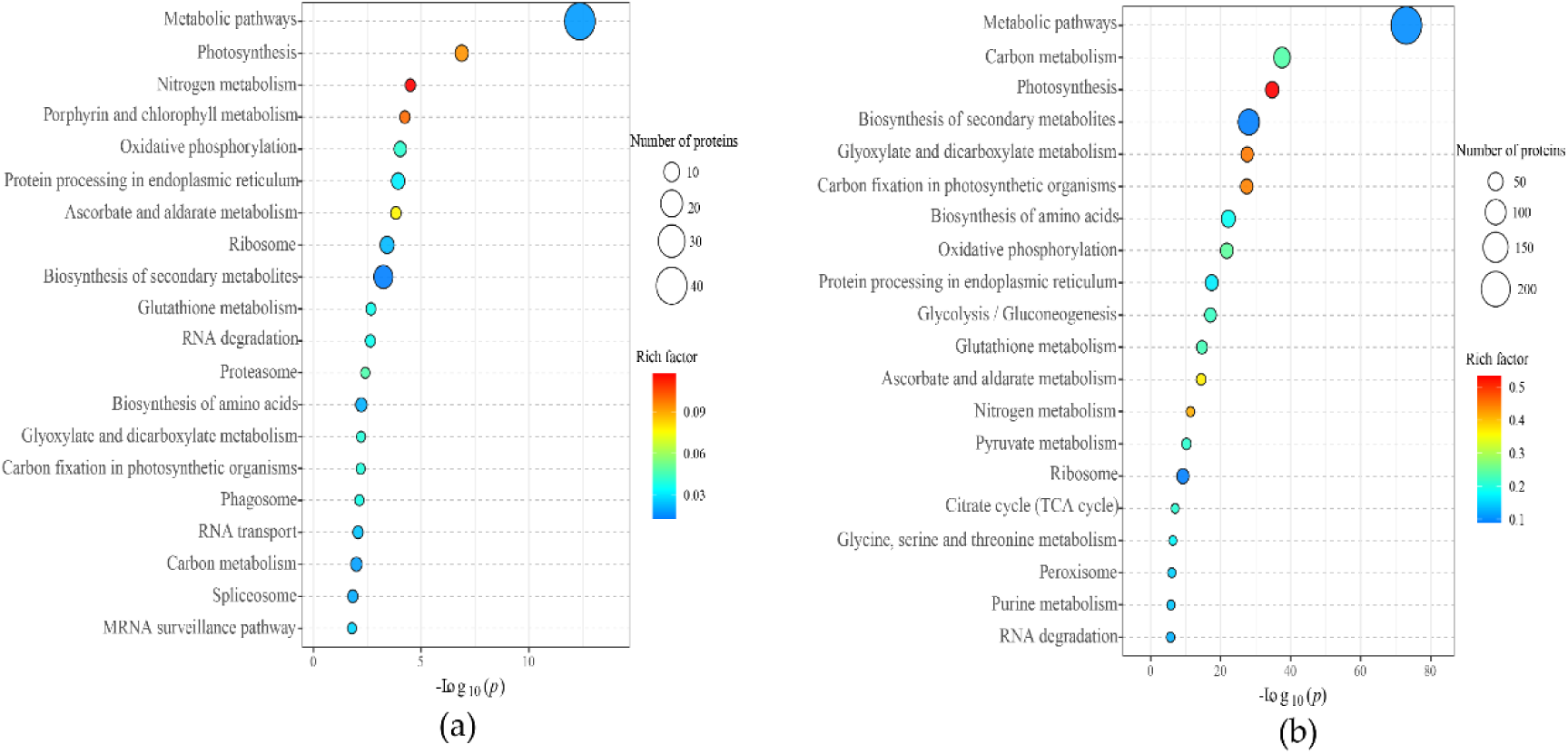
Kyoto Encyclopedia of Genes and Genome (KEGG) pathway enrichment bubble map of the top 20 DEPs. (a) KEGG pathway enrichment map of the top 20 DEPs of the GD vs. GW group. (b) KEGG pathway enrichment map of the top 20 DEPs of the XD vs. XW group. The vertical coordinate indicates the enriched pathway, while the horizontal coordinate indicates the negative logarithmic transformation of the p value. A p value of <0.05 indicates that the function is significantly enriched, with smaller p values indicating significantly higher functional enrichment; the color of the circle denotes the rich factor: Rich factor = (A/B)/(C/D), where A denotes the num-ber of differential proteins annotated to the term, B denotes the total number of differential pro-teins annotated to the term, C denotes the number of background proteins of the term, and D de-notes the total number of background proteins of the term. A larger Rich factor indicates a higher level of enrichment, and the size of the circle indicates the number of DEPs enriched in the path-way.

The KEGG enrichment analysis of the 20 DEPs in the MEpink module indicated that they were enriched in 12 and 3 KEGG metabolic pathways in the G and X varieties, respectively, under drought stress. Among them, DEPs of variety G were significantly enriched in five KEGG pathways (*p* < 0.05) including mRNA surveillance pathway, ribosome, phenylalanine metabolism, photosynthesis, and phagosome, while DEPs of variety X were significantly enriched in three KEGG pathways (*p* < 0.05) including photosynthesis, phagosome, and plant–pathogen interactions. Thus, the KEGG enrichment pathways of DEPs differed between the two different drought-resistant varieties, implying that they may have different metabolic pathways in response to drought stress.

### Protein division in the MEpink module

In the MEpink module, there were 17 DEPs in the variety G under drought stress. All of them were upregulated, except for two DEPs shared with the variety X (one is a chaperone protein DnaJ [ID: M8A0F5] and the other is an uncharacterized protein [ID: A0A446J984]), and the remaining DEPs were majorly involved in energy metabolism, protein translation, RNA processing, amino acid metabolism, etc. These DEPs were 9-kDa polypeptide (ID: A0A0Q3IET4), carbamoyl-phosphate synthase (ID: Q9AXS0), temperature stress-induced apolipoproteins (ID: I1IBR5), photosystem I subunit VII (ID: A0A446S4S2), PAP_fibrillin domain-containing protein (ID: A0A446MFG9), SpoU_sub_bind domain- containing protein (ID: A0A0Q3FXR3), SRP54 domain-containing protein (ID: A0A446KTE5), isoleucyl-tRNA synthetase (ID : I1H766), and seven uncharacterized proteins (ID: A0A446SZM6, A0A453AN53, A0A510B3S2, A0A446QWZ0, M5BPX4, A0A3B6FW66, and A0A453N829). The variety X had three DEPs (excluding the two DEPs shared with the variety G) under drought stress, all of which were upregulated in expression; these were peptide chain release factor (ID: M8CEY9) and two uncharacterized proteins (ID: A0A3B5YS83 and A0A3B6A287) and were primarily involved in RNA processing, protein stabilization, plant photosynthesis, intracellular signal transduction, etc.

## Discussion

### Common pathways of coping with drought stress in different drought-tolerant varieties

DnaJ protein is a heat stress protein isolated from Escherichia coli by Georgopoulos et al. in 1980 [32]; it is primarily involved in protein folding, translocation, secretion, and localization and denatured protein reversion and degradation and plays a vital role in maintaining intracellular protein homeostasis under adverse conditions and the stability of protein complexes [33]. By studying DnaJ protein expression in pepper placenta, Fan found that DnaJ protein may be involved in the plant response to abiotic stress during the biosynthesis of stimulatory compounds [34]. Wang investigated the function of tomato (Lycopersicon esculentum) chloroplast-targeted DnaJ protein (LeCDJ2) and its ectopic expression in transgenic tobacco and found that salicylic acid, drought, and pathogen attack induced the expression of LeCDJ2 and that the ectopic expression of LeCDJ2 in transgenic tobacco decreased the production of superoxide anion radical (O^2−^) and hydrogen peroxide (H_2_O_2_) under drought stress; moreover, the overexpression of tomato chloroplast-targeted DnaJ improved drought and cyanobacterial resistance of transgenic tobacco [35]. In the present study, DnaJ protein expression was significantly upregulated in the two different oat varieties, suggesting that DnaJ protein may be involved in the response to drought stress, which is similar to the findings reported by Rodrigues and Walia [36,37].

### Different pathways of coping with drought stress in different drought-tolerant varieties

Photosystem I (PSI) is a multisubunit membrane complex that catalyzes the electron transfer of iron redox proteins that bind to FNR and decrease NADP+. The PSI complex in higher plants has 4–80-kDa polypeptides in addition to LHCI, of which the 9-kDa polypeptide subunit is a protease that may function when plants are grown under unsuitable conditions [38]. In this study, the expression of the 9-kDa polypeptide was significantly upregulated in the strongly drought-resistant variety G, whereas its expression in the water-sensitive variety X was insignificant, suggesting that strongly drought-resistant oat varieties may respond to drought stress by improving energy production during cellular photosynthesis, thereby re-establishing homeostasis and complex energy metabolic pathways in plants, which is consistent with a previous discovery that the expression of proteins involved in photosynthesis is upregulated in drought-resistant maize varieties [39].

Spermidine is a precursor for the production of compounds that act as second messengers, such as nitric oxide (NO) and polyamines (polyamines include spermidine, spermidine, and putrescine), involved in the regulation of stress resistance in plants [40–43]. For example, NO and H^2^O^2^ are critical molecules in regulating abscisic acid (ABA)-induced stomatal closure in plants [44]; exogenous chitosan and spermine attenuate drought-induced oxidative damage and increase the content of protective metabolites total phenols and flavonoids in white clover (Trifolium repens L.) [43]. The subunit of carbamoyl-phosphate synthase is a catalytic enzyme necessary for the conversion of ornithine to citrulline during arginine biosynthesis and is involved in plant resistance reactions [45]. This was further verified using the results of the present study, in which this enzyme was significantly upregulated in expression under drought stress in drought-resistant variety, catalyzing the synthesis of ornithine into citrulline and releasing energy for continued plant growth under adverse conditions [43], but not in water-sensitive variety.

Lipoproteins are critical enzymes of the lutein cycle and play an essential role in protecting the photosynthetic apparatus of plants from oxidative damage caused by excessive light exposure [46]. The two homologous cDNAs from wheat (TaTIL) and Arabidopsis (AtTIL) encode lipoproteins containing three structurally conserved regions that are possible ligands with diverse structures and functions for the synthesis of phytosteroid hormones, such as 2,4-epi-oligolactone, which enhance plant tolerance to heat and cold stress [47,48]. Attil lipid transport proteins are also involved in regulating plant tolerance to oxidative stress [49]. Plant temperature-induced lipid (TIL) transport protein counteracts severe heat stress-induced lipid peroxidation [50]. AtTIL, a lipid-like transport protein gene, plays an essential role in plant salt tolerance [51]. However, research on TIL concerning drought resistance in plants has not been reported. In this study, TIL transport proteins were significantly upregulated in the drought-tolerant variety under drought stress conditions. In contrast, the expression abundance did not change remarkably in the water-sensitive variety, revealing that TIL transport proteins may be involved in regulating the drought tolerance of plants and improving their drought resistance.

Plastids are organelles that store organic matter and perform photosynthesis in plants and are generally divided into chloroplasts, colored bodies, and white bodies. There are plastoglobules with a diameter of 30 nm to 5 μm in plastids, which are lipoprotein particles [52]. A family of fibrillin (FBN) was found in this lipoprotein. The FBN gene was induced by stresses, such as high temperature, low temperature, drought, salt, herbicides, and strong light, mediating the resistance signaling of ABA and regulating the synthesis of jasmonic acid, triacylglycerol, and plastoquinone [53–56]. In this study, the expression of PAP_fibrillin domain-containing protein in the variety G was significantly upregulated under drought stress, which was consistent with the increased expression of FBN protein under drought stress in Arabidopsis, tomato, potato, tobacco, and other plants in previous studies [53,57,58]. This shows that the PAP_fibrillin domain-containing protein may be involved in the recognition and response of oat chloroplasts to drought stress, and further investigation is required to determine whether this protein plays a significant role in the recognition and response to drought stress and how it does so.

Selective splicing in plants is a vital post-transcriptional regulatory mechanism that regulates gene expression and ultimately affects plant morphology and function [59]. Serine/arginine-rich proteins are mainly involved in the assembly and splicing of eukaryotic mRNA precursors. They are critical factors in the assembly and splicing of mRNA spliceosomes, playing a unique regulatory role in their splicing mechanisms [60]. It was found that plants can respond to adversity stress through the regulation of splicing factors; for example, variable splicing of the OsWRKY45 allele is involved in drought stress response [61], and the overexpression of ScMYBAS1-2 and ScMYBAS1-3 splice transcripts promotes plant growth changes under water insufficiency and drought conditions [62]. In this study, SRP54 domain-containing protein was significantly upregulated under drought stress in the variety G but not in the variety X . These results suggest that SRP54 domain-containing protein may play an important role in mediating oat plants’ response to drought stress as Shearing factor.

In the MEpink module in this study, seven uncharacterized proteins were significantly upregulated under drought stress conditions in the variety G but not in the variety X. These proteins with unknown structure and function may play essential roles in the regulatory network of drought stress and improve drought resistance in oats. Therefore, further studies on these uncharacterized proteins will help elucidate the molecular mechanisms of oat response to drought stress.

## Conciusions

Our study explored the mechanisms of drought tolerance in two different drought-tolerant oat varieties from phenotypic, physiological and proteomic perspectives, and clarified the interconnections between the physiological and proteomic factors contributing to it. In conclusion, our findings contribute to further understanding of drought tolerance mechanisms in different drought-tolerant oat varieties.

### Author Contributions

CC and MB designed and performed the experiments; CC, MB and YZ performed experiments together; MB and YZ analyzed the data, prepared figures and wrote the paper; XW and WL revised the manuscript; All authors read and approved the final manuscript.

## Acknowledgments

This work was funded by the National Modern Agricultural Industry Technology System of China (CARS-34); We also thank Gaolong Yin at Shanghai Bioprofile Technology Company Ltd. for his technical support in proteomics.

## Conflicts of Interest

The authors have no conflict of interest.

## Notes

### Competing Interest Statement

The authors have declared no competing interest.

